# Genomic duplications shaped current retrotransposon position distribution in human

**DOI:** 10.1101/819284

**Authors:** Andrea Riba, Maria Rita Fumagalli, Michele Caselle, Matteo Osella

## Abstract

Retrotransposons, DNA sequences capable of creating copies of themselves, compose about half of the human genome and played a central role in the evolution of mammals. Their current position in the host genome is the result of the retrotranscription process and of the following host genome evolution. We apply a model from statistical physics to show that the genomic distribution of the two most populated classes of retrotransposons in human deviates from random placement, and that this deviation increases with time. The time dependence suggests a major role of the host genome dynamics in shaping the current retrotransposon distributions. In fact, these distributions can be explained by a stochastic process of random placement due to retrotranscription followed by genome expansion and sequence duplications. Besides the inherent interest in understanding the origin of current retrotransposon distributions, this work sets a general analytical framework to analyze quantitatively the effects of genome evolutionary dynamics on the distribution of genomic elements.

## I. INTRODUCTION

Transposable elements (TEs or transposons) are sequences of DNA able to move within a host genome. Transposons are a crucial force driving genome evolution, they are found in all the genomes, with very few exceptions, and compose nearly half of the human genome [1–4].

A large number of TEs in human belongs to the class of retrotransposons (REs), which proliferates through a “copy-and-paste” mechanism. Indeed, they are first transcribed into RNA intermediates, and then reverse transcribed into the host genome at a new position [3–5]. The two most abundant RE classes in human are SINEs (Short Interspersed Nuclear Elements) and LINEs (Long Interspersed Nuclear Elements) whose main representative members are Alu and L1 families, respectively [3, 6]. These two families globally consist of ≈2 millions elements and together account for nearly the 30% of our genome [1, 7]. Both Alus and L1s can then be divided into subfamilies depending on the nucleotide sequence of the active TEs driving the subfamily expansion [3, 5]. Throughout the paper, the term *family* refers to Alu and L1, while the term *subfamily* refers to these subgroups.

The proliferation dynamics of these subfamilies has a relatively short timescale. In fact, after a rapid burst of amplifications and insertions, during which every new element becomes a potential source of retrotranpositions, the subfamily turns silent or inactive [6, 8]. Further rounds of transcription and insertion of the REs are typically prevented by the accumulation of sequence mutations, rearrangements, truncations or specific methylations able to inactivate the process [2, 3, 6, 8]. Thus, the REs that can be identified in the current genomes are generally a fossil track of the history of subsequent birth-extinction cycles of different mobile sequences, with few subfamilies still currently expanding in the human genome, such as the L1H subfamily [2, 9].

Therefore, the genomic distributions of RE subfamilies reflect possible specific preferences or biases of the insertion mechanism during the subfamily active period, but carry also information about the most relevant evolutionary forces driving the rearrangement of the host genome after the subfamily expansion. For example, evolutionary moves such as genome expansion or duplications of DNA segments should alter the RE distributions in specific ways.

This work addresses the evolutionary mechanisms that have shaped the current distributions of genomic distances between REs of different subfamilies, focusing on members of the abundant Alu and L1 families as relevant examples. Despite the well recognized importance of retrotransposons in the evolution of genomes, several aspects of their proliferation dynamics are still obscure. The RE position on the genome is arguably the simplest observable that contains information about this dynamics, and nonetheless has still to be fully characterized and explained.

We will show, using analytical arguments and data analysis, that these empirical distributions can be explained as a result of a process of insertion in random genomic positions, followed by sequence duplications and expansion of the host genome. A model based on these mechanisms not only can explain empirical RE distributions but also naturally leads to predictions (e.g., on the role of RE density) that were confirmed by data analysis.

Besides the interest that the still partial understanding of the REs dynamics and its interactions with the host genome has in itself [7, 10–12], the theoretical framework developed in this paper is general and can be easily extended to the study of spatial distributions of other functional genomic elements along the genome.

## II. MATERIALS AND METHODS

### A. Genomic data

The human genome sequence (assembly GRCh38/hg38) was downloaded from UCSC database [13]. We considered only sequences referred to chromosomes 1-22, X, Y. The number and genomic positions of transposable elements in hg38 were downloaded from RepeatMasker official website [14] (*RepeatMaskeropen* − 4.0.5 - Repeat Library 20140131). The analysis were performed on Alu and L1 subfamilies with more than 1000 elements, in order to guarantee sufficient statistics. We included in the analysis a total of 32 Alu subfamilies and 107 L1 subfamilies. We verified that the typical size of Alu elements is around 300 bps, while of L1s is less than 1 kbp (see Supplementary Figure S1) [1, 11]. The distance between successive REs of each subfamily was calculated as the difference between the start genomic coordinate of an element and the stop coordinate of the previous one. We verified that an alternative definition of the inter-REs distance using half-length coordinate of REs does not alter our conclusions, as it was expected given that our analysis is based on large scale observations.

The distances between the start (end) of each chromosome and the first (last) retrotransposable element of the considered subfamily have been excluded. Distances falling in centromeric and pericentromeric regions were also neglected since these regions are usually highly repetitive, rich in copy number variations (CNVs) and difficult to sequence properly. As a consequences, they contain few extremely long inter-RE distances that shows in the distributions as outliers of the order of few Mbp.

At the end of this filtering process less than 50 inter-RE distances have been discarded for each subfamily and the effective human genome length has been reduced to about 2.8 Gbp, depending on the subfamily. Moreover, also REs inserted in the middle of pre-existing elements of the same subfamily were discarted from the analysis to avoid an excess of zeros in the distance distribution.

This procedure is necessary to ensure the validity of our assumption of point-like REs and affects only about 3% of L1 elements while is completely negligible (i.e., < 0.01%) for the Alu family.

### B. Stick-breaking process as a null model for random positioning of small genomic elements

We consider a set of *B* genomic elements whose length is small enough relatively to the genome or chromosome size *L* of interest. In this case, we can introduce a formal analogy with the stick-breaking (SB) process or the fragmentation process well studied in statistical physics [15–18]. This process considers point-like fractures randomly positioned on a stick or polymer of fixed length. The length distribution of the fragments after *B* breaks are positioned can be analyzed analytically. To make the analogy exact, the genomic elements of interest have to be well approximated by point-like elements, as it is the case for REs given their short length with respect to the genome or to chromosomes [1, 11].

The analytical solution of the SB process can be used to test if the genomic elements of interest are indeed randomly placed along a genomic sequence. According to ref. [15], the expected number of inter-break distances equal to *x* after the random placement of *B* breaks on a support of length *L* is

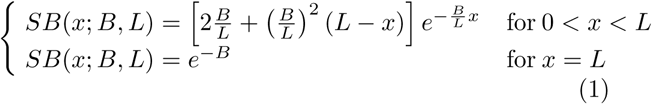

The probability distribution *p*_*SB*_(*x*; *B, L*) is simply obtained by normalizing *SB*(*x*; *B, L*) with the total number of distances *B* + 1.

### C. A measure of deviation from random placement

We developed a measure of the deviation of an empirical distribution of inter-element distances from the parameter-free distribution expected for random placement and described by Eq. 1. This measure is proportional to the area between the empirical and null model distributions, in analogy with the Cramér-von Mises criterion [19]. More specifically, it is the integral between the two cumulative distributions. However, since the mean and standard deviation of a SB depend on the density of elements (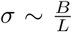 for large *B*), we normalize this integral by the density of each subfamily in order to make the deviations comparable for sets with a different number of elements. As discussed in more detail in the SI, distributions relative to random placement for different values of *B* and *L* collapse to a single functional form thanks to the normalization. Thus, the distance from the expected SB in Eq. (1) for a given subfamily *i* can be defined as

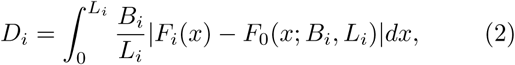

where *F*_*i*_ and *F*_0_ are the empirical and theoretical cumulative distributions. The normalization factor *B*_*i*_*/L*_*i*_ is the density of subfamily *i* and allows the comparison of *D*_*i*_ for subfamilies of different abundances.

### D. The insertion-elimination model

This section formalizes a model describing the dynamics of distances between a set of point-like elements (REs in our case) on a genome under the hypothesis that the two main evolutionary forces are genome expansion due to insertion of other genomic elements, and random deletions of the point-like elements under consideration, which effectively leads to the “fusion” of two existing distances. The simplest assumption is that the probability of insertion of new sequences is uniform over the whole genome and that all elements have the same probability of being deleted. These are precisely the assumptions considered in ref. [20]. In this case, the probability that a new insertion event expands the distance *x* between two existing REs is proportional to *x*. If we introduce a typical length scale *λ* of the inserted sequences and the rate *γ* at which on average an insertion happens, we can describe the process as

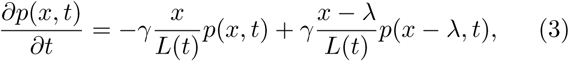

where, for the moment, we are neglecting the elimination of REs. The equation (3) describes the time evolution of the distribution *p*(*x, t*) of distances of length *x* while the support *L*(*t*) is expanding. It assumes that the probability of inserting a sequence of length *λ* into an existing inter-element distance of length *x* is simply proportional to its length (i.e., to 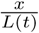), and that there is an approximately constant rate *γ* of new insertions. As described in more detail in the Supporting Information, Eq. 3 can be solved in the continuous limit that is valid as long as the distances are long enough as it is always the case for REs. The solution is simply 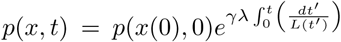, where *p*(*x*(0), 0) is the distance distribution before the dynamics starts.

The initial condition is given by the stick-breaking process with *B*_0_ breaks described in the previous section since we are assuming an initial random placement. In other words, *p*(*x*(0), 0) = *p*_*SB*_(*x*(0); *B*_0_, *L*(0)) from Eq. 1. As shown in more detail in the Supporting Information, the factor multiplying the stick-breaking initial condition in the solution is just a rescaling of the support. In fact, the evolved inter-RE distances at time *t* is simply described by

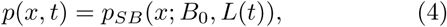

where *L*(*t*) = *L*(0) + *γλt*. In conclusion, the shape and the functional form of the initial SB distribution are not modified by the expansion dynamics, and the distribution can be still described by a SB on an expanded genome *L*(*t*). The model can be generalized by assuming that there is a distribution *ρ*(*λ*) of the lengths of insertions, but the result does not change qualitatively. Analogously, deletions of genomic segments can be considered, but also in this case the distribution would not change its shape as long as the deletions are randomly placed.

So far we implicitly assumed that the inserted sequences have a length *λ* smaller than existing inter-RE distances. If this is not always the case (i.e., if there are *x*_0_ < *λ*), the last term in Eq. 3 is not relevant for these short distances, and the solution has an additional term of the form

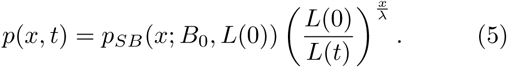

Therefore, the complete solution has two terms: for short distances (*x* < *λ*) there is an exponential behaviour that deviates from the SB distribution, while for long distances the process is described by an expanded SB. This behavior is confirmed by simulations as explained in detail in the Results section.

The introduction of RE elimination, for example due to sequence mutations, would not alter the picture above. In fact, if we take the solution of the SB process with *B* breaks in Eq. 1 and we randomly eliminate a certain number *n* of breaks, the resulting distribution would still be a solution of a SB process with *B* − *n* breaks. This intuitive result is confirmed by numerical simulations in Fig. 3.

**FIG. 1:**
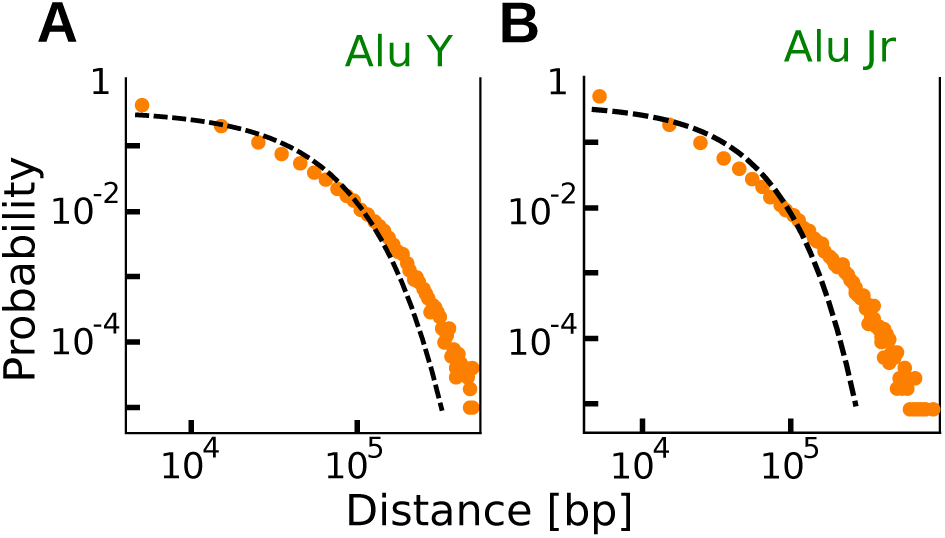
The genomic distribution of retrotransposable elements shows deviations from random placement. The Figure shows the empirical inter-REs distance distribution (symbols) of two different Alu subfamilies in the human genome. Specifically, panels A and B refer to genomic distribution of Alu Jr and Alu Y subfamilies. The dashed black line represents the parameter-free analytical expectation given by the null model based on a random-placement hypothesis (Eq. (1)). More examples supporting similar deviations from random placement can be found in the Supplementary Figure S2.

**FIG. 2:**
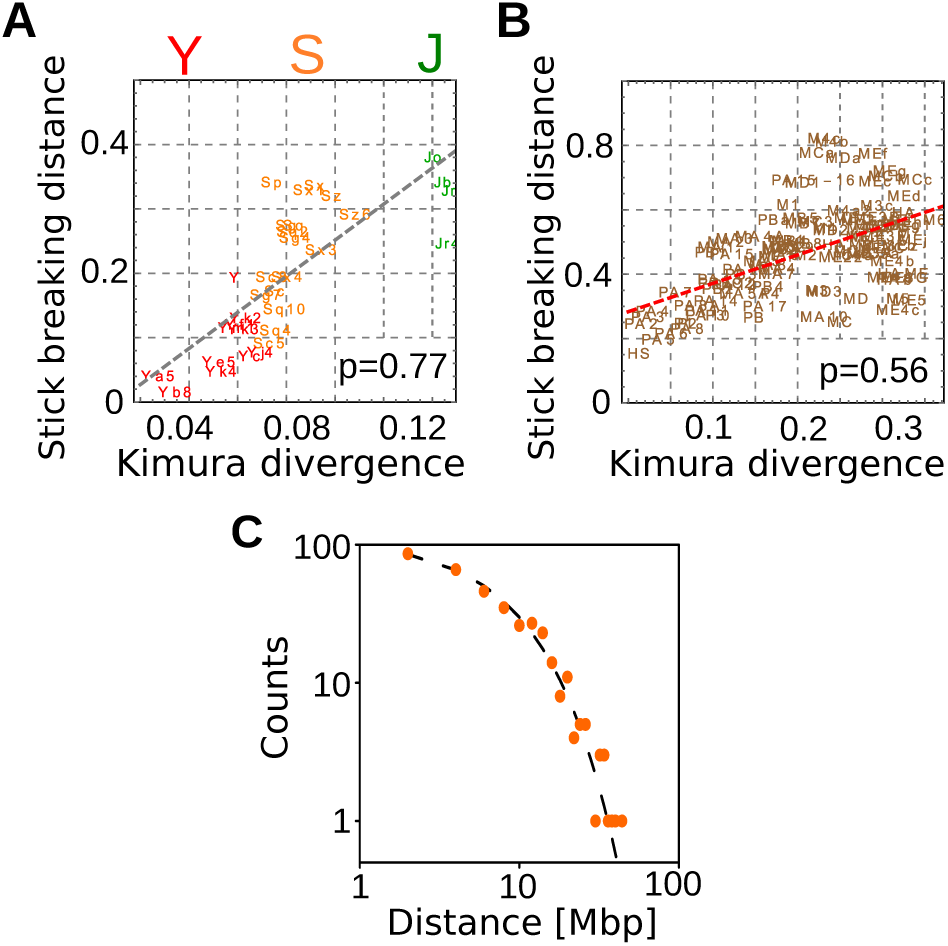
The deviation from random placement increases with the RE subfamily age. The deviations (estimated with Eq. (2)) between the empirical inter-RE distance distributions and the corresponding expectation for random placement are shown as a function of the age of retrotrasposon subfamilies for Alus (panel A) and L1 (panel B). The clear correlation is supported by the correlation coefficients reported in the figures while the dashed lines are linear fits. The age of different subfamilies is estimated by the Kimura divergence corrected for CpG hypermutability (more detail in the SI). Panel C shows that very recently inserted retrotransposons are distributed along the genome in perfect agreement with the model based on random insertion. Specifically, we considered 367 L1 elements detected in a sample of 25 individual human genomes (data from ref. [9]) and not included in the reference genome.

**FIG. 3:**
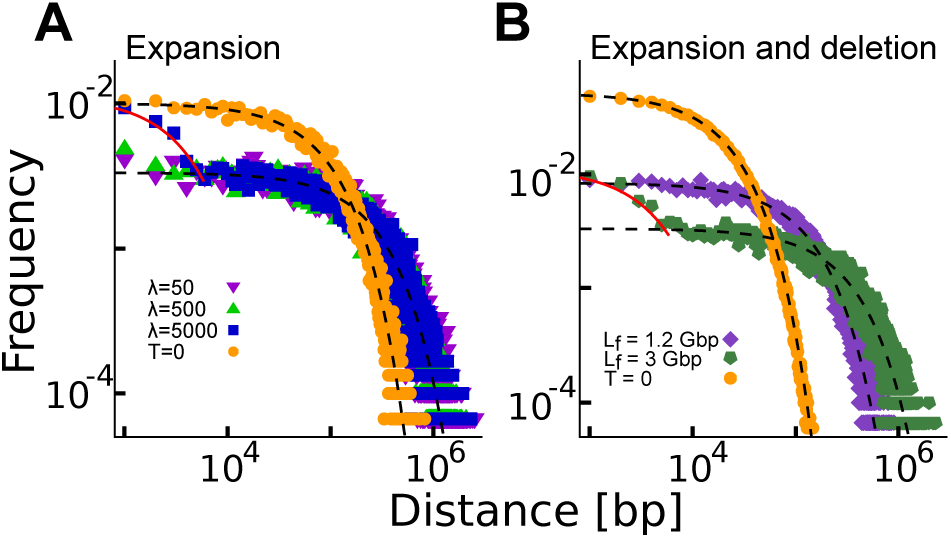
Genome expansion and loss of REs cannot affect the shape of the inter-RE distribution. Panel A shows the simulated distribution of distances between *B*_0_ = 10^4^ points randomly placed on a segment of length *L*_0_ = 10^9^ (orange dots) and the effect of an expansion process due to insertions of segments of different lengths *λ* represented by the different symbols described in the legend. The final genome size is M*L*_*f*_ = 3 · 10^9^ for all the distributions. In panel B we added in the simulations the random elimination of “break” points. The initial distribution of distances (orange dots) is given by *B*_0_ = 5 · 10^4^ points randomly placed on a segment of length *L*_0_ = 10^9^. After genome expansion (with *λ* = 5000) and RE random elimination at different rates (different symbols), we report the inter-RE distance distributions when the final number of REs has reached *B*_*f*_ = 10^4^. Black dashed lines correspond to the null model distribution with the correct number of REs *B* and genome size *L* as in Eq. (4). Continuous red lines represent the solution for *x* < *λ* in Eq. (5). Only for large genome expansion driven by large insertions a small deviation from random placement can be observed.

### E. A model including genomic duplications

The model presented in the previous section can be effectively extended in order to take into account the result of genomic duplications. The extension is based on the observation that a duplication event that also duplicates some of REs of interest adds to the distance distribution precisely the distances between the duplicated elements. Specifically, the distribution *p*(*x, t*) of inter-RE distances will have some new distances of a certain length *x* that depends on the relative position of the REs that have been duplicated. This effect can be phenomenologically captured in the model by assuming that there is an external source of new distances described by a term *q*(*x*). This term represents the probability of adding a distance of size *x* as a result of a duplication event and its form should depend on the existing distance distribution at the moment the duplication occurs. We can also introduce a parameter *µ* for the rate of duplications.

Therefore, a model that includes both genome expansion and duplication of genome portions can be formalized by the equation

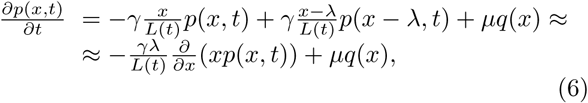

which simply extends Eq. 3 by adding the source term *µq*(*x*). In the continuous limit, the method of characteristic leads to the following system of equations

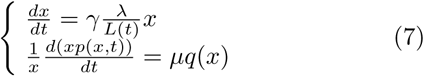

Rewriting Eq. (7) and assuming a linear increase of the genome size *L*(*t*) = *L*(0) + *φt* (where *φ* is the combined growth of the genome given by expansion and insertions) we can derive

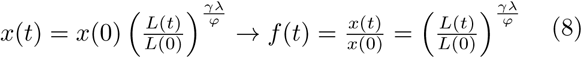

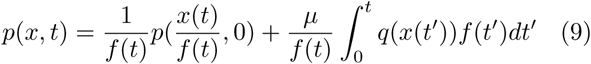

The solution (9) is composed of two terms: the first one describes the expansion of the original SB while the second represent the source expansion over time. *f*(*t*) is a monotonic and increasing function of time, as defined in (8), that weights the initial condition relative to the source at a specified time *t*. It is obtained from the first equation in (7). Note that 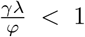, since *φ* includes both expansion (*γ*) and source (*µ*). By substituting *p*_*SB*_(*x*; *B*_0_, *L*(0)) as initial condition for the first term on the right in Eq. 9, we still find a SB

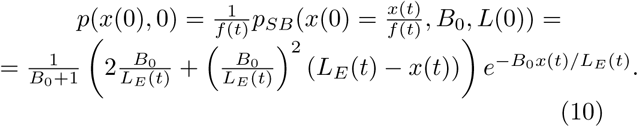

The expanded genome is now *L*_*E*_(*t*) = *L*(0)*f* (*t*), since the whole genome *L*(*t*) contains also the contribution of the (expanded) source. In the limit *φ* → *γλ* we recover the result of the former section. In order to fully solve the expansion-insertion model we have to choose an explicit form for the source function *q*(*x*). While its precise functional form is in principle unknown, several considerations (see Results), point to a fast decreasing function of *x* (such as an exponential decay).

### F. Direct estimate of the effect of duplications on inter-RE distances

If the source term *q*(*x*) is a fastly decreasing function of *x*, such as an exponential decay, the probability of adding long distances to the inter-break distribution is extremely small. Therefore, the presence of a source term in our dynamical model is not expected to alter significantly the shape of the right tail of the initial inter-break distribution. In our case, the initial distribution is supposed to be the stick-breaking solution in Eq. (1). Moreover, we have previously shown that the expansion of the support by random sequence insertion cannot change the functional form of the initial random distribution, Eqs. (9) and (10). Relying on these considerations, the right-tail of the inter-RE distribution should be well approximated by random placement of a smaller number of elements with the correct normalization and length of the support, respectively *B*_0_ and *L*_*E*_(*t*) in Eq. (10). More formally, normalizing Eq. (9) by the total number of breaks and taking into account the relation in Eq. (10), the current distribution can decompose as follow

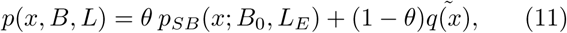

where *θ* is the fraction of RE distances that are still distributed according to the initial SB solution. The effective source 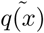 is proportional to the source *q*(*x*) weighted over the time with *f* (*t*), i.e., the initial source expanded and averaged over *f* (*t*).

Within the assumption that the source *q*(*x*) of new segments only affects the short scale distances, we can assume that the long-distance tail of the distribution of *B* breaks on a genome *L* can be well explained by a stick-breaking solution with a smaller initial number of breaks *B*_0_, while the short-distance region of the distribution is a superposition of this stick-breaking process with the contribution of duplications captured by the fastly decaying source term 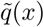. This also implies the existence of a minimum distance *x*_*min*_, above which the effect of the external source is negligible (Supplementary Figure S6). In other words, for *x* > *x*_*min*_ the observed distances should be well fitted by a SB solution with an initial number of breaks *B*_0_ (smaller than the empirical subfamily size), while for *x* < *x*_*min*_ we have a superposition of the SB solution and of the source term 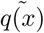. Following the same idea from Clauset et al. [21], we identify *x*_*min*_, *B* and *L* for each subfamily using a maximum likelihood approach to find the best possible fit of Eq. (11). With this approach we can directly estimate both the initial number of breaks *B*_0_, and the fraction of duplicated Res 1 − *θ*. Finally, the source term 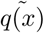 can be deduced as the difference between the empirical distribution and the best estimate of *p*_*SB*_(*x*; *B*_0_, *L*_*E*_).

The procedure has been applied to all Alu subfamilies. The estimated thresholds *x*_*min*_ are of the order of 10^5^ bp. The resulting source terms 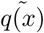 can be generally well fitted by an exponentially decreasing function (Supplementary Figure S7) and their averages correlate with the density *B/L* of the subfamily as discussed in the Results section and shown in detail in the SI. For the older Alu subfamilies, 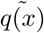 can be better approximated with a double exponential function (Figure 4B).

**FIG. 4:**
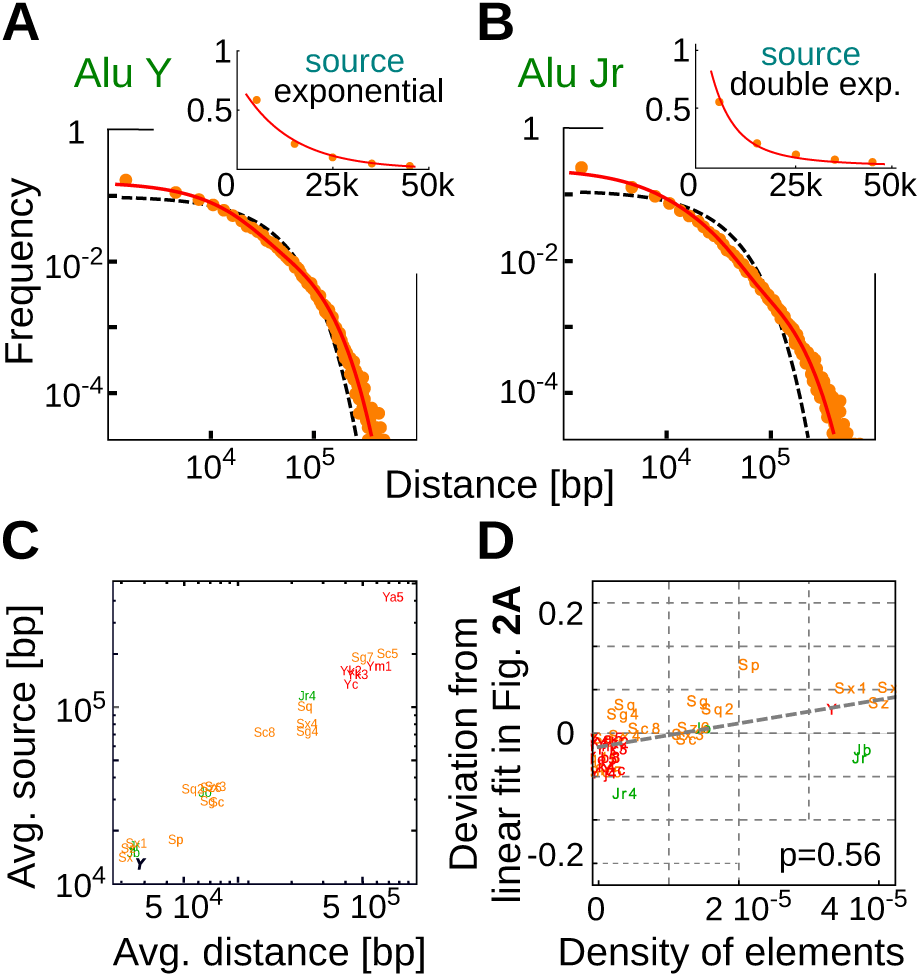
A model including local duplications and genome expansion reproduces the empirical distances between retrotransposable elements. Panel A and B show the inter-Alu distance distributions (red dots) for two subfamilies of different age. The best fit with a superposition of a stick-breaking solution and a source term capturing the effect of duplications (Eq. (9)) is plotted as a continuous orange line. The expectation for random positioning (Eq. (1)) is also reported as dashed black line for reference. The insets show that the source terms (red dots) estimated with the methodology explained in the Methods section can be well approximated by an exponential function for the relatively young AluY subfamily (A) and by a double exponential function for the older AluJr subfamily (B). Panel C reports the correlation between the average length of the added distances by the duplication process (average of the source term in Eq. (11)) and the average distances between REs of the different subfamilies. Finally, panel D shows that the deviation from random positioning is dependent on the subfamily density and on the subfamily age. In fact, the deviation from the linear dependence on age reported in Figure 2A can be, at least partially, explained by the differences in subfamily densities.

## III. RESULTS

### A. Retrotransposons are not randomly distributed along the genome

The first question we address in this section is how far the empirical RE distributions are from the simplest assumption of random genomic placement. An eventual deviation from random placement can be due to biases in the insertion process itself during the activity of the sub-family, such as specific sequence preferences for insertion, as well as from subsequent neutral or selective processes of the genome such as rearrangements, duplications or sequence specific RE deletions. The presence of biases in the insertion mechanism is still debated. While there is convincing evidence that the insertion process of REs actually occurs at random positions along the genome [3, 9, 22], specific sequence preferences for the insertion sites have also been reported [23]. On the other hand, several works highlighted non-random properties of the current RE positioning that could in principle be ascribed to subsequent genomic processes. For example, there is a density enrichment of specific subfamilies in genomic regions with high or low GC content [1, 24, 25], and a signal of formation of clusters of REs [26]. On a global scale, the distributions of distances between REs of different families has been observed to deviate from random positioning by visually comparing the empirical distributions with randomly generated surrogate datasets of RE positions [20].

A mathematical model for random placement would allow us to place the above observations in a well defined quantitative setting and to actually measure possible deviations from a random placement assumption. This model can be easily formulated by realizing an analogy between the positioning of relatively small genomic elements and a well studied process in statistical physics, i.e., the stick-breaking process (SB). The SB process was originally formulated as a model of the stochastic fragmentation of a polymer chain [15, 16, 27, 28] and it is described in detail in the Methods section. If the breaks are assumed to be completely random, the SB provides an analytical expression for the expected distribution of fragment lengths that only depends on the length of the polymer *L* and the number of breaks *B* (Eq. (1) in the Methods section). The analogy with the RE insertion process is based on the observation that the REs can be safely considered point-like, since their length is extremely small with respect to the genome itself [1, 11], and thus are equivalent to the point breaks in the SB formulation. In fact, the average RE sequence length is around 300 bp for the Alu class and 1 kbp for the L1 elements (see Supplementary Figure S1), which are negligible with respect to the genome length in human (around 3.2 Gbp). Therefore, the distance distribution between RE elements should be precisely equivalent to the length distribution of fragments defined by the SB process if the REs are randomly positioned in the genome.

The two parameters *B* and *L* that define the distance distribution (Eq. (1)) for REs simply correspond to the number of retrotransposons of a specific subfamily and to the genome length considered. Figure 1 and Supplementary Figure S2 show the comparison between the SB parameter-free predictions and some illustrative examples of empirical inter-RE distance distributions. This comparison clearly shows evidence of non-random positioning. The empirical deviations from a SB are due to an enrichment of both short and long distances, suggesting that the mechanisms that shaped current RE distance distributions must have both increased the “clustering” of retrotransposons and correspondingly fostered the presence of very distant elements. The same trend is observed if the inter-RE distributions are analyzed on single chromosomes rather than on the whole genome (Supplementary Figure S2). Even if the deviation from random placement makes the empirical distribution more “long-tailed” with respect to the null expectation, there is no clear evidence of a power-law behaviour of these distributions as was previously suggested [20]. In the following, the analytical model will be used to disentangle if these deviations are mainly due to specific biases in the insertion mechanism itself or to the genomic rearrangements after the subfamily inactivation.

Several previous analysis reported that different families can have specific preferences for genomic regions with different GC content at the level of initial insertions or because of subsequent sequence-specific selection of RE elements [1, 24–26]. For example, both Alus and L1s have been reported to have an insertion preference for AT-rich regions [1, 25], even though current distributions can show an opposite bias: high density of Alus in GC-rich regions, and viceversa a high density of L1s in GC-poor regions [25]. In order to test if the deviations from random positioning we observe are simply driven by GC content, we divided the genome in GC rich and GC poor regions, and analyze the inter-RE distance distributions limited to these regions for different subfamilies. The details of this procedure are reported in the SI. Even though the density of REs is indeed dependent on the GC content, the deviations of the inter-RE distance distributions from the random expectation do not differ in genomic regions with different GC content (Supplementary Figure S3). Therefore, the general trend reported in Fig. 1 of an accumulation of short RE distances is robust with respect to the GC content, suggesting that it cannot be simply explained by a random-positioning process with different insertion probabilities depending on the GC content.

### B. The age of a retrotransposon subfamily is strongly correlated with its deviation from random placement on the genome

The deviations from random positioning described in the previous section can be dominated by biases in the insertion process or by subsequent genome evolution mechanisms that can continuously reshape the RE distributions even after their inactivation. In the latter case, the deviation should be more apparent for old subfamilies. In fact, the inter-RE distributions of young or currently active subfamilies should be mainly determined by the choice of retrotranscription sites. Therefore, a time dependence of the deviations from random positioning would suggest that these are not mainly driven by the insertion process but rather by subsequent time-dependent processes. In order to test this, we introduce a measure that quantifies the deviations of empirical inter-RE distance distributions from the corresponding distributions predicted by the SB process. This measure is analogous to the Cramér-von Mises criterion [19], and it is based on the area between the two cumulative distributions (empirical and theoretical), normalized by the RE density (Eq. (2), see Methods for more details). The normalization is necessary in order to safely compare deviations for RE subfamilies that have different global densities.

Figure 2A and B shows that this distance from random positioning is well correlated with the age of the RE subfamilies both for Alus (Pearson correlation p=0.77) and for L1s (p=0.56). The age of a subfamily can be estimated by evaluating the number of mutations between the RE sequences and a reference sequence [29]. The reference sequences for each subfamily come from Repbase [30] and the Kimura divergences are automatically inferred by RepeatMasker [14]. While Figure 2 reports the Kimura divergence as the estimate of the subfamily age, the trend is conserved if other estimates, such as the Jukes-Cantor divergence, are used. The clear correlation suggests that the process that drove the positioning of RE is time dependent and that the retrotransposition sites are instead more close to random placement as testified by the fact that recent subfamilies are better described by the null model. As a further test, we analyzed the inter-RE distances for 367 L1H elements detected in a sample of 25 individual human genomes (data from ref. [9]). These insertions are not fixed in the human population since they are not present in the reference genome. Therefore, we can confidently assume that this set of L1s is originated by very recent retrotransposition events, and thus genomic rearrangements did not have the time to reshape the RE positions. Fig. 2C shows that indeed their relative distances are perfectly compatible with the random expectation, supporting previous evidence that retrotransposition occurs in random positions [22, 26, 31, 32]

Also in this case, we tested that the trend is consistent if GC-rich or GC-poor genomic regions are considered separately (see Supplementary Figure S4).

### C. Genome expansion and sequence mutations cannot explain current retrotransposon position distributions

The previous section strongly supports a scenario in which random insertion of REs has been followed by rearrangements of the host genome that reshaped their positions. Now, the question is which specific genomic events may explain the features of current RE distributions. A previous analysis suggested that genome expansion due to random insertions of new genomic elements coupled with progressive elimination of REs (e.g., by mutation-induced “degradation” of their sequences) could explain current spatial RE distributions [20]. However, this section will show that a model based on these two simple mechanisms, called insertion-elimination model (IE) by the authors [20], cannot actually fully explain the empirical RE distributions.

First, most of the insertions driving genome expansion are actually due to transposable elements themselves. As we discussed previously, the length of these elements is less than ∼1 kbp (Supplementary Figure S1), thus typically much shorter than the inter-RE distances. If we consider a genome expansion driven by the insertion of small elements, we can show analytically that the shape of the RE inter-distance distribution does not change. The only effect of genome expansion is to rescale all distances by the same factor as they simply expand with the same rate of the genome itself. The analytical proof of this intuitive behavior is reported in the Methods section. The master equation in Eq. (3) is a good approximation of the process for insertion lengths *λ* ≲ *L/B* and the solution of this equation is still the solution of a SB process but on a longer support (Eq. (4)). Analogously, the elimination of REs cannot change the functional form of their inter-distance distribution. If the RE are simply eliminated at random (as it would be the case for random mutations), the resulting distribution of their relative distances would still be random, only with a smaller number of REs.

Only in presence of a large number of inserted sequences that are much longer than existing inter-RE distances we can observe a deviation from a random distribution (Fig. 3 and the Methods section). In this case, the “preferential-attachment” mechanism suggested by Sellis et al. [20] can take place. Short distances with respect to *λ* cannot be created by expansion of even shorter distances. Additionally, they are progressively less likely to be hit by a new insertion since the insertion is proportional to the segment length. The overall effect is that the final distribution is an overlap of two distributions corresponding to two different rates of expansion. Simulations suggest that this effect becomes relevant only after an extremely high number of insertion events (e.g., at least doubling the initial genome size) as reported in Fig. 3. This large number of insertions of very long sequences sounds very unlikely as an explanation of the empirical deviations from random placement of REs. As a further test we simulated for a couple of illustrative RE subfamilies, a realistic genome expansion by considering the insertion of other REs from younger subfamilies and sequence duplications (not involving the REs under analysis) directly estimated from the human genome sequence (Supplementary Figure S5). Also in this case, the empirical deviations from random placement cannot be explained by the model.

The addition of elimination of REs in the process does not change the results. Intuitively, random elimination of breaks simply decreases the parameter *B*_0_ in the stick-breaking process without affecting the shape of the distribution. Therefore, the combination of genome expansion and RE elimination would still lead to a distribution equivalent to the one obtained by considering genome expansion alone. We tested with numerical simulations (Fig. 3B) that indeed the effect of genome expansion and random loss of REs cannot generate the significant deviations from random placement empirically observed in Fig. 1. In summary, we can state that an insertion-elimination process is unlikely to be the main driver of current RE distribution in the genome. The next section will show how adding sequence duplications in the evolutionary model can actually explain the empirical inter-RE distributions.

### D. Genomic duplication is the key ingredient to explain the observed distributions of retrotransposons

The previous section showed that simple models based on genome expansion by random insertions and RE sequence mutations are not enough to explain the empirical distributions of REs. A main evolutionary force of genome evolution that we have not considered so far is sequence duplication. Genomic duplication is a major source of genomic rearrangements and it is quite common across the whole phylogenetic tree [33, 34].

A duplication of a random genomic sequence of a given length can either contain a certain number of the REs under analysis or none of them. If the duplicated segment does not contain any of the REs under study, we are back to the model of the previous section. In fact, no additional “break” is added, and the net effect of the duplication is just a sequence insertion in a given position that expands a certain inter-RE distance. On the other hand, if the duplicated segment does contain some REs, the relative distances between the duplicated REs will add to the distance distribution. This can be modeled by assuming that duplications effectively represent a source of new REs and thus of new distances, as detailed more formally in the Methods section. This source term essentially captures the probability of adding a inter-RE distance of given length as a result of a duplication event. It phenomenologically contains unknown and complex information on the length distribution of the duplicated sequences and on the probability of capturing a certain number of REs with a certain relative distance distribution inside the duplicated segment. Although it is extremely hard to characterize the precise functional form of this source term, we can devise reasonable approximations and test them a posteriori.

If the REs are initially inserted at random on the genome, as supported by the results in the previous sections, the distribution of the REs on a sequence that is duplicated is expected to be still a random distribution described by the stick-breaking process (Eq. (4)) but on a support of size given by the length of the duplicated region. Therefore, a duplication event adds to the initial random distribution another random distribution but defined on a segment of much smaller length. This means that the probability of adding an inter-RE distance of length *x* by a duplication event is well approximated by an exponentially decreasing function of *x* (Eq. (1)). Therefore, as long as the process has not yet significantly changed the initial RE distribution, the source term should be well described by an exponential function. In fact, the solution of our model with this assumption for the source term (reported in the Methods section Eq. (9)) can fit very well the empirical distributions of relatively young subfamilies. An illustrative example is reported in Fig. 4A.

On the other hand, if the dynamics had a sufficiently long time to alter the initial random distribution through duplications, the inter-RE distances that are added by a duplication event will not be well described by an exponential function anymore. After several duplication events, a duplicated segment will contain inter-RE distances following a distribution that already has an increased number of relatively short distances, thus generating a positive feedback that drives an effective strong clustering of REs. As a consequence, the source term should be better described by a decay faster than exponential. This is indeed the case for older subfamilies. The model with a double exponential functional form for the source term is able to fit much better the empirical distributions of older subfamilies as shown in Figure 4B for one example.

In any case, duplications are expected to generate an excess of short distances (or more clustering) with respect to random placement. Therefore, the right tail of the inter-RE distance distribution should not be significantly affected by the duplication process. This observation allows us to devise a simple method for estimating the source term directly from data. The hypothesis is that the right part of the distribution should be well described by random placement of a number of REs *B*_0_ smaller than the one currently observed *B*. This corresponds to the initial distribution of the REs of the subfamily we are studying. The subsequent duplications are expected to increase the number of REs (from *B*_0_ to the observed value *B*) and only change significantly the left part of the distribution. Therefore, we can assume that current distributions are a superposition of a stick-breaking process which dominates for large distances and of a term defined by the source that defines the short-distance part of the distribution (Supplementary Figure S6). As explained more formally in the Methods section, under this assumptions we can directly estimate the source term and the initial number of REs *B*_0_ with a simple fitting procedure. The results of this direct estimate of the source terms confirm the above considerations: an exponential source term (compatible with a SB solution) is enough to explain the distance distributions of most subfamilies while for the older Alu subfamilies a steeper function, such as a double exponential, better explains the data (Supplementary Figure S7). Moreover, the fraction of duplicated REs (*B* −*B*_0_)*/B* is correlated with the age of the RE subfamily as expected in a scenario of subsequent duplications after a random retrotranscription process (Supplementary Figure S8). The estimated percentage of REs that were duplicated can be as high as ∼ 85% for the older Alu J subfamilies, while for the youngest Alu Ya5 is approximately the 8% (Supplementary Table 1).

A natural consequence of the considerations above is an expected correlation between the average inter-RE distance added by a duplication event (i.e., the average of the source term) and the density of a subfamily. In fact, a duplicated segment simply contains a smaller part of the RE distribution of a given subfamily that is well approximated by a SB at the beginning. The more elements are present, the shorter is their typical distance and thus the shorter will also be the typical RE distances duplicated in the evolutionary process. Figure 4C shows that this prediction is confirmed by empirical data. The average inter-RE distance is well correlated with the average distances added by the source term.

A more subtle prediction of our model with genomic duplications concerns the role of RE density. We have shown that the deviation from random placement is correlated with the time elapsed from the birth of a subfamily. However, if this deviation is mainly driven by duplications, subfamilies of similar age but with different densities on the genome should display different degrees of deviations. A random duplication event is likely to include a number REs of a subfamily that depends on their density on the genome. Therefore, the variability that can be observed in Fig. 2A and B should be explained by a variability in subfamilies densities. We tested this prediction by measuring the deviation from the linear fit in Fig. 2 as a function of RE density for the different subfamilies. The results are reported in Fig. 4D and confirms the presence of a correlation. In other words, the deviation from random placement depends on the age of a subfamily but also on its density on the genome, further supporting the hypothesis that random duplications were a main evolutionary force in shaping current inter-RE distance distributions.

## IV. DISCUSSION

Retrotransposable elements, and in particular L1s and Alus, compose a large fraction of the human genome and their role in genome evolution has been increasingly recognized [33, 35–37]. Transposable elements impact the genome in a variety of ways since they can promote structural rearrangements, contain regulatory elements, harbor transcription and splicing sites, and are involved in the production of non-coding RNAs [7, 11, 12, 38–40]. A large number of genetic diseases and cancers have been linked to mobile elements [5, 41–43], although the causal relation is still unclear [44–47]. Despite their importance, a clear understanding of their dynamics in the genome is still elusive.

This work focuses on the position distribution of retro-transposons at the genome scale, and on the role of the host genome dynamics in shaping their relative genomic distances. To this aim, we introduced a formal analogy between the retrotransposition process and the well studied process in statistical physics of random insertion of breaks in a polymer chain [15, 17, 28, 48]. Leveraging on this analogy, we could rephrase in a quantitative setting several longstanding questions. First, we proved that current positions of most RE subfamilies are not randomly distributed along the genome. While previous studies made this observation [20], we could assessed it quantitatively and, more importantly, define a natural measure of the extent of the deviation from random placement of empirical distributions. This measure indicated that the degree of non-randomness of RE positions is strongly correlated with the age of the subfamily in analysis. More specifically, the position of REs of very recent or still active subfamilies is well described by random placement, while this description becomes progressively less accurate as the age of the subfamily increases. Specifically, REs tend to become more clustered over time than expected from the random model.

Therefore, the analysis of recent or active subfamilies further confirms that retrotranscription occurs approximately at random sites in the genome (at least at this large observation scale), giving a quantitative support to previous empirical observations [3, 9, 22]. Note that “local constraints” for fixation of retrotransposition events could be present and induce specific biases in the insertion sites. For example, sequence preferences linked to GC content were suggested for L1s [23]. In this regard, we analyzed in detail the role of GC content to show that the phenomenology of RE progressive clustering here described do not change qualitatively in regions with different GC content. There is indeed a dependence of RE density on GC content, but we showed that this difference is not the driver of the observed inter-RE distance distributions. Analogously, insertions in coding genes or in regulatory regions can be detrimental, and thus under strong negative selection. However, our analysis suggests that these constraints do not play a major role in the RE positioning at the genomic or at the chromosomal scale. Indeed, only a small fraction of our genome is actually coding, and a recent estimate based on mutational load considerations of the functional fraction of our genome leads to a conservative upper bound of 25% [49]. As previously suggested, most transposon insertions seem indeed to be neutral or only mildly deleterious and thus simply subjected to genetic drift [50]

However, the inter-RE distance distribution for most subfamilies is far from random. As the time dependence of this non-randomness suggests, the progressive evolution of the host genome must have reshaped the RE positioning in a specific way. While the genome evolves and rearrange, the RE already present will be passively moved and repositioned in the genome. We thus tried to pinpoint the main evolutionary mechanisms responsible for the specific non-random features of current RE distributions by testing different evolutionary models. We first analysed a model based on genome expansion and RE elimination that was previously proposed as a candidate to explain RE positions [20]. However, analytical calculations and extensive simulations showed that these mechanisms are not sufficient to quantitatively explain the empirical distributions. Therefore, we added genomic duplications in the model, and the resulting effect on the distribution of inter-RE distance was precisely the one empirically observed: a time-depenent increase of the REs at short distances that could well match the data relative to different subfamilies. Several tests of the model, such as the effect of RE density on the typical distance between duplicated REs or on the deviation from random position, further confirmed that genomic duplications and genome expansion are the key ingredients to explain current retrotransposon positions in human. It would be extremely interesting to apply this analysis to other species to test the generality of this result.

A major role for genomic duplication in moulding current genomes has been widely recognized since the pioneering work of Ohno [51]. Therfore, it should not be surprising that current RE distributions have been strongly influenced by this evolutionary mechanism as our study demonstrates. However, the rough estimates that our method provides for the fraction of duplicated REs range from few percent for recent familes, to more than half for older ones. The fact that such a large fraction of old transposable elements is likely to come from duplications rather than direct retrotrasposition is puzzling. While this could be an overestimation of our method, it should be noted that duplications due to non-allelic homologous recombination, such as large segmental duplications [33, 52], can be promoted by the presence of repeated sequences such as transposons. For example, Alus are often found at the border of a particular class of segmental duplications called tandem duplication [33, 53]. Therefore, the presence of retrotransposons can enhance the duplication probability. This interplay would establish a positive feedback between duplication and retrotransposon density that indeed will drive the inter-RE distance distribution toward the “clustering” we observe, and could explain the large fraction of duplicated RE we estimate for old subfamilies.

Finally, the analytical framework developed here thanks to the analogy with the stick-breaking process can be a powerful tool. In fact, it gives analytic and parameter-free predictions for random positioning of small genomic elements that can be directly compared with the empirical ones. In the same framework, we introduced a measure of non-randomness and developed simple but tractable evolutionary models that can be used to quantify and disentangle the different evolutionary contributions to the positioning of the genomic elements. Given its generality, this approach can be naturally extended to the study of other elements such as, for example, small regulatory sequences or single nucleotide polymorphisms.

## V. ACKNOWLEDGEMENTS

We thank Marco Cosentino Lagomarsino and Mattia Furlan for their critical reading of the manuscript. This work was supported by the “Departments of Excellence 20182022” Grant awarded by the Italian Ministry of Education, University and Research (MIUR) (Grant No. L. 232/2016). The work of MRF was supported by AIRC under IG2018-ID 21558 project-449 PI Pusch Michael.

