## Supplementary Information for "Genomic duplications shaped current retrotransposon position distribution in human"

October 23, 2019

#### S1 Kimura divergence

Different models are available to measure the evolutionary divergence between two DNA sequences. Insertion time of a retrotransposable element (RE) can be determined as the ratio between the divergence $d$ of the RE from a consensus sequence and the total substitution rate $r$ Ray et al. [2008].

Substitution rate indicates the probability of mutation of a generic base to another one ([A, C, G, T]). The overall substitution rate in human is estimated to be of the order of $2 \cdot 10^{-8}$ mutation per nucleotide per generation Keightley [2012], Nachman and Crowell [2000]. Hence, assuming a generation time of $\approx 15$ years the mutation rate results $\approx 1.3 \cdot 10^{-9}$ mutations per nucleotide per year.

Kimura model (Kimura [1980], Nei and Kumar [2000]) allows to estimate for the divergence taking into account two different mutational processes. It has been noticed that it is easier to mutate between nucleotide with the same chemistry, so that mutation between two purines or pyrimidines (i.e. $A \leftrightarrow G$ and $C \leftrightarrow T$) have a transitions rate ($\alpha$) larger than other mutations (transversions rate, $\beta$ e.g. $A \leftrightarrow C$). Thus, Kimura total substitution rate is $r = \alpha + 2\beta$ and the divergence is

$$d = -\frac{1}{2}\ln(1 - 2p_1 - p_2) - \frac{1}{4}\ln(1 - 2p_2) \tag{1}$$

where $p_1$ and $p_2$ are the fraction of transitions and transversions, respectively.

Further corrections can be added accounting for the hypermutability of CpG dinucleotides (Hodgkinson and Eyre-Walker [2011], Liu et al. [2009], Duret and Arndt [2008]). RepeatMasker implements this correction neglecting the mutated CpG dinucleotides. Indeed, Alus have an high GC content and the estimate of their age is highly influenced by the CpG content correction.

#### S2 Distance from SB distribution

Defining a distance between data and the null model can give a quantitative estimate of the observed deviations. As in the main text, we define the distance for a given subfamily $i$ between the distribution of inter-RE distances and the stick-breaking as

$$D_{SB} = \int_0^{L_i} \frac{B_i}{L_i} |F_i(x) - F_0(x; B_i, L_i)| dx \,. \tag{2}$$

The first term $F_i(x)$ is the cumulative distribution of the empirical inter-REs distances, while $F_0(x; B_i, L_i) = \left[1 - \left(1 - x\frac{B_i}{L_i}\frac{1}{B_i+1}\right) e^{-xB_i/L_i}\right]$ is the cumulative distribution of the corresponding null model (SB).

The cumulative stick-breaking distribution can be expressed using the rescaled variable $y = xB_i/L_i$. It is important to note the standard deviation of a SB distribution is approximated for large $B$ to $\frac{L}{B}$. In other words, the rescaling collapses the expected SB distributions to a distribution with the same average and the same standard deviation for all sub-families.

Since the quantity $\frac{x}{L}$ is expected to be very small compared to $B$ (for our datasets it is usually $\frac{x}{L} < 10^{-3}$.), we can neglect the linear term $\frac{y}{B_i+1}$. In this case, Eq.(2) can be approximated as:

$$D_{SB} \approx \int_0^{B_i} |F_i(y) - \left(1 - e^{-y}\right)| dy \tag{3}$$

The above expression shows that the distances we calculated can be compared between different subfamilies. Indeed, we are measuring the distance of different empirical distributions from almost the same function $\approx 1 - e^{-y}$. Note that the upper limit of the integral, that depends on the specific subfamily $i$, is never reached. In fact $y = B_i$ would imply $x = L_i$.

#### S3 GC-rich and GC-poor regions

We define *GC content* of a DNA sequence the ratio between the total number of bases and the number of Cs and Gs present in the sequence. The GC content is calculated over non-overlapping windows of $10^6 bp$, excluding those with more than 20% of masked bases. These windows should be sufficiently large to contain elements even for the smaller subfamilies. A window is considered GC-rich when its GC content is $> 41\%$ Bernardi [2015]. Then, contiguous windows with similar GC content are collapsed in single regions and we obtained a total of 521 different regions, 263 defined as GC-rich and 258 as GC-poor. This procedure guarantees a sufficient “minimal” length of the regions in order to have a significant statistic of REs elements inside them. The same procedure was repeated on the masked sequence of the human genome where the repeats are masked by capital Ns UCS. In this case we neglected windows with more than 50% of masked bases. The results obtained with the two methods are comparable (not shown).

#### S4 Expansion-deletion model

Under the hypothesis of randomly distributed events the probability of insertion or deletion in a specific position is uniform over the whole genome. As a consequence the probability that an expansion/deletion event modifies an inter-REs distance $x$ is proportional to $x$. It is possible to write a very general equation for the evolution of the distances distribution:

$$\begin{aligned} \frac{\partial p(x,t)}{\partial t} = & \gamma_e \frac{x-\lambda_e}{L(t)} p(x-\lambda_e,t) - \gamma_e \frac{x}{L(t)} p(x,t) \\ & + \gamma_d \frac{x+\lambda_d}{L(t)} p(x+\lambda_d,t) - \gamma_d \frac{x}{L(t)} p(x,t) \end{aligned} \tag{4}$$

accounting for the increase and decrease in size of preexisting distances due to insertions and deletions of length $\lambda_e$ and $\lambda_d$ respectively. The factors $\gamma_e$ and $\gamma_d$ are the rate of expansion/deletion. In the limit of "small" $\lambda_d$ and $\lambda_e$ (essentially, $<< L/B$), Eq. (4) can be approximated introducing partial derivatives

$$\begin{aligned} \frac{\partial p(x,t)}{\partial t} = & -\gamma_e \frac{\lambda_e}{L(t)} \frac{\partial (x p(x,t))}{\partial x} \\ & + \gamma_d \frac{\lambda_d}{L(t)} \frac{\partial (x p(x,t))}{\partial x} \end{aligned} \tag{5}$$

and it assumes the simple form:

$$\frac{\partial p(x,t)}{\partial t} = -\gamma \frac{\lambda}{L(t)} \frac{\partial (x p(x,t))}{\partial x} \tag{6}$$

In this case, the factor $\gamma\lambda = \gamma_e\lambda_e - \gamma_d\lambda_d$ represents the effective expansion/deletion parameter. Equation (6) can be solved using the method of characteristic:

$$\begin{cases} \frac{dt}{dk} = 1 \\ \frac{dx}{dk} = \gamma \frac{\lambda}{L(k)} x \\ \frac{dp(k)}{dk} = -\gamma \frac{\lambda}{L(k)} p(k) \end{cases} \tag{7}$$

Identifying the variable $k$ with the time and integrating we obtain:

$$\begin{cases} x(t) = x(0) e^{\gamma\lambda \int_0^t \frac{dt'}{L(t')}} \\ p(x(t),t) = p(x(0),0)\, e^{-\int_0^t \gamma \frac{\lambda}{L(t')} dt'} \end{cases} \tag{8}$$

The evolved distribution encompasses a rescaling over time of the variable $x$, by the factor $e^{-\gamma\lambda \int_0^t \frac{dt'}{L(t')}}$. This rescaling can also be seen as the total genome lenght growing over time, because of the new insertions. Indeed, imposing as initial condition

$$\begin{aligned} & p(x(0),0) = p_{SB}(x(0); B_0, L(0)) = \\ & \frac{1}{B_0+1} \left( 2\frac{B_0}{L(0)} + \left(\frac{B_0}{L(0)}\right)^2 (L(0) - x(0)) \right) e^{-\frac{B_0}{L(0)} x(0)} + e^{-B_0} \delta\left(x(0) - L(0)\right), \end{aligned} \tag{9}$$

the expanded distribution results:

$$p(x,t) = \frac{1}{B_0+1} \left[ \left( 2\frac{B_0}{L(t)} + \left(\frac{B_0}{L(t)}\right)^2 (L(t) - x) \right) e^{-\frac{B_0}{L(t)} x} + e^{-B_0} \delta\left(x - L(t)\right) \right] \tag{10}$$

where $L(t) = L(0)e^{\gamma\lambda \int_0^t \frac{}{L(t')} dt'}$. The total number of REs is constant during the process, and the normalization condition $\int_0^{L(t)} p(x,t)dx = 1$ must hold and as a consequence

$$p(x,t) = p_{SB}\left(x;\ B_0, L(t)\right). \tag{11}$$

The expansion process does not modify the shape of the distribution. Moreover, we can derive an explicit solution for the genome size as a function of time

$$L(t) = L(0)e^{\gamma\lambda \int_0^t \frac{dt'}{L(t')}} \to L(t) = L(0) + \gamma\lambda t \tag{12}$$

Since the insertion rate $\gamma$ is kept constant, the linear increase of $L$ in time is consistent. The above solution is valid for positive and negative values of $\lambda\gamma$ (i.e. expansion term in Eq. (5) dominates over deletion or vice versa). In the second case both $x$ and $L$ decrease in time. Note that we should impose some limit to the increase (decrease) of genome size. However, we are interested in modeling a "realistic" expansion process spanning a finite time. Thus, we can consider $L(t)$ and $L(0)$ to be of the same order of magnitude.

For short inter-REs distances (i.e. $\lambda > x$) the discrete equation (4) should be modified and its continuous approximation is no longer valid. In this region the distribution evolves according to:

$$\frac{\partial p(x,t)}{\partial t} = -\gamma \frac{x}{L(t)} p(x,t) \tag{13}$$

Imposing $L(t)$ as in Eq. (12), the evolved distribution at time $t$ results:

$$\begin{aligned} p(x,t) &= p(x,0)e^{-\gamma x \int \frac{dt}{L(t)}} \\ &= p_{SB}(x; B_0, L(0)) \left(\frac{L(0)}{L(t)}\right)^{\frac{x}{\lambda}}. \end{aligned} \tag{14}$$

The complete solution should be given by a superimposition of Eq. (11) and Eq. (14).

### S5 Duplication source

The solution of a pure expansion model presented above, can be extended in order to account the presence of an external source of new inter-REs distances

$$\frac{\partial p(x,t)}{\partial t} = -\gamma \frac{\lambda}{L(t)} \frac{\partial \left(x p(x,t)\right)}{\partial x} + \mu q(x,t) \tag{15}$$

Using the method of characteristic and rewriting Eq. (15) as a total derivative of the function $xp(x,t)$

$$\begin{cases} \frac{dx}{dt} = \gamma \frac{\lambda}{L(t)} x \\ \frac{1}{x} \frac{d(xp(x,t))}{dt} = \mu q(x) \end{cases} \tag{16}$$

By assuming a linear increase of the genome size $L(t) = L(0) + \varphi t$, where $\varphi$ is a combined function of expansion and insertion contributions, the solution becomes

$$\begin{cases} \frac{x(t)}{x(0)} = \left(\frac{L(t)}{L(0)}\right)^{\frac{\gamma\lambda}{\varphi}} = f(t) \\ p(x,t) = \frac{1}{f(t)} p(\frac{x}{f(t)}, 0) + \frac{\mu}{f(t)} \int_0^t q(x(t')) f(t') dt' \end{cases} \quad (17)$$

Substituting $p_{SB}(x_0; B_0, L_0)$ as initial condition the first term on the right in Eq. (17) is still a stick-breaking function:

$$\frac{1}{f(t)} p_{SB}(\frac{x}{f(t)}; B_0, L_0) \;\; = \;\; \frac{1}{B_0+1} \left( 2\frac{B_0}{L_E(t)} + \left(\frac{B_0}{L_E(t)}\right)^2 (L_E(t) - x) \right) e^{-B_0 x / L_E(t)}. \quad (18)$$

The expanded genome is now $L_E(t) = L(0)f(t)$, since the whole genome $L(t)$ contains also the contribution of the (expanded) source. In the limit $\varphi \to \gamma\lambda$ we recover the result of the former section. In order to fully solve the expansion-insertion model we have to choose an explicit form for the source function $q(x)$.

Segmental duplication is a relevant mechanism for genome evolution. The duplication of a DNA sequence can imply the passive duplication of the retrotransposons eventually contained in the sequence. The source term $q(x,t)$ can be seen as the *effective* source of inter-REs distances resulting from these events. In the main text $q(x)$ with an exponential shape has been choosen as source term for the fits in Figure 4.

### S6 Maximum likelihood estimate

In the main text, we suggested that the tail of the inter-REs distance distribution could be described by the stick-breaking function, with the correct parameters. We consider the probability distribution

$$\left\{ p(x, x_{min}, B_0, L_E) = \frac{1}{Z} \left( 2\frac{B_0}{L_E} + \left(\frac{B_0}{L_E}\right)^2 (L_E - x) \right) e^{-\frac{B_0}{L_E} x} \right. \quad (19)$$

where the factor $Z$ guarantees the normalization of the stick-breaking function on the range $[x_{min}, L_E]$ and $L_E$ indicates the expanded genome.

Unfortunately, you cannot apply the maximum likelihood criterion in an analytical way to such distribution. Then we applied the following numerical procedure: for a fixed threshold $x_{min}$, we use a two step numerical procedure to pinpoint the number of breaks $B_0$ and the genome $L_E$.

We repeat the estimate procedure of the parameters for different values of the threshold $x_{min}$ and we selected the value of $x_{min}$ (and as consequence, the corresponding optimal couple $\{B_0, L_E\}$) that minimize the Kolmogorov-Smirnoff distance between the stick-breaking function and the data. The procedure described was applied to all the 32 selected Alus subfamilies. For seven small subfamilies it was impossible to estimate the parameters (usually, we obtained $x_{min} = 0$ and $B_0 > B_{real}$).

For most of the other subfamilies $x_{min} \approx 10^5$. The estimated sources are reported in Fig.S7.

#### S7 Supplementary Figures

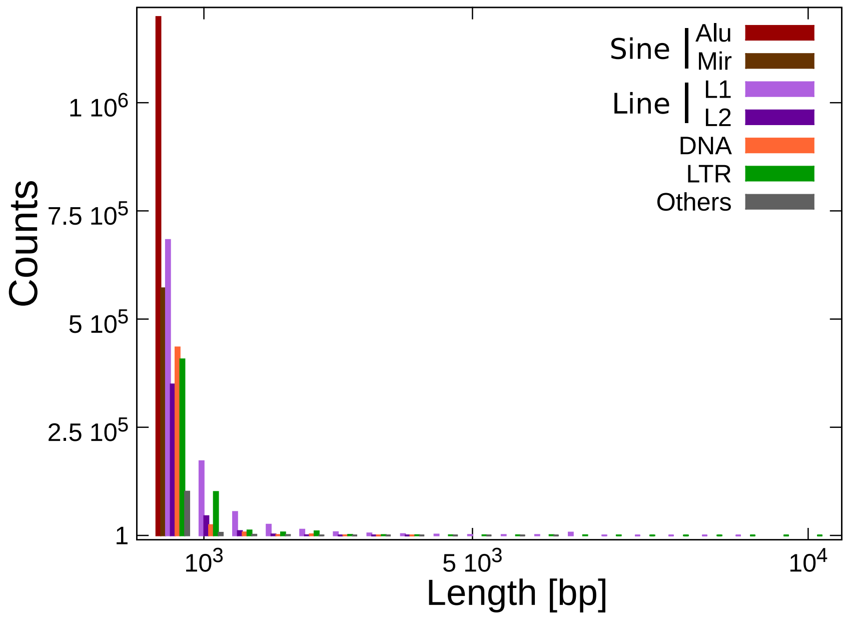

**Supplementary Figure S1. Length distribution of transposable elements in human genome.** Most of the transposable element insertions in the human genome are less than 1 kbp long, including LINEs elements. Satellite repeats are neglected. Bin size is 500 bp.

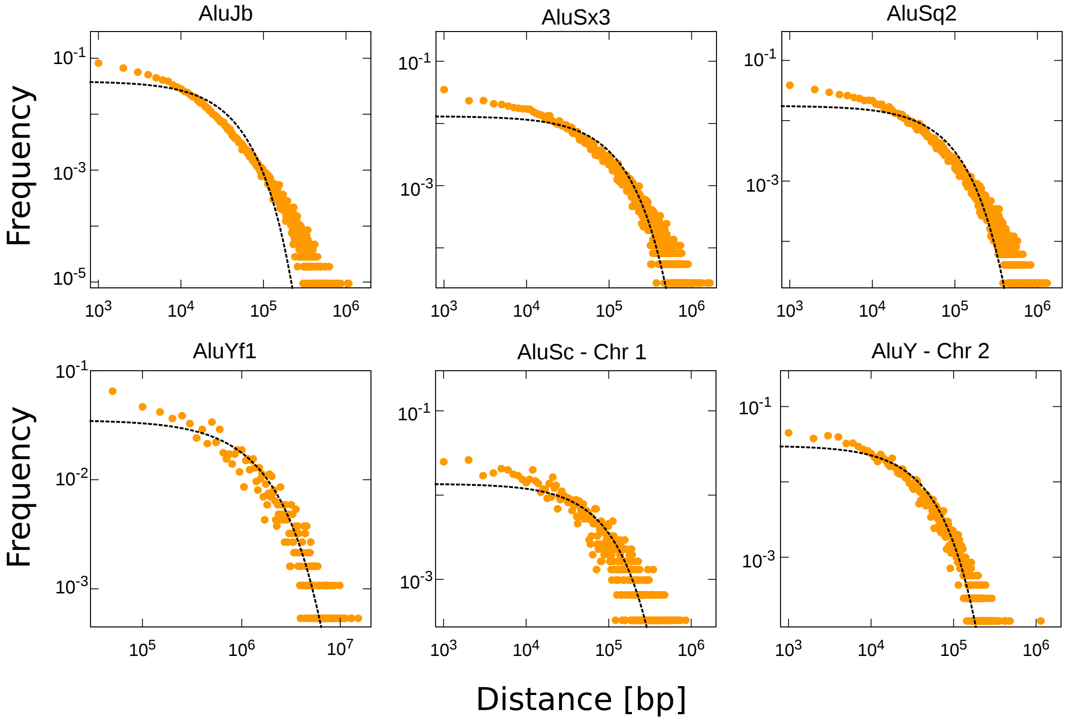

**Supplementary Figure S2. Examples of inter-Alus distances distribution.** The Figure shows few examples of inter-Alus distance distribution (orange dots) for different subfamilies. If chromosome is not specified, data refer to the whole genome distribution. The predictions from the stick-breaking process (Eq. 1 of the main text) relative to each subfamily are shown as black dashed lines.

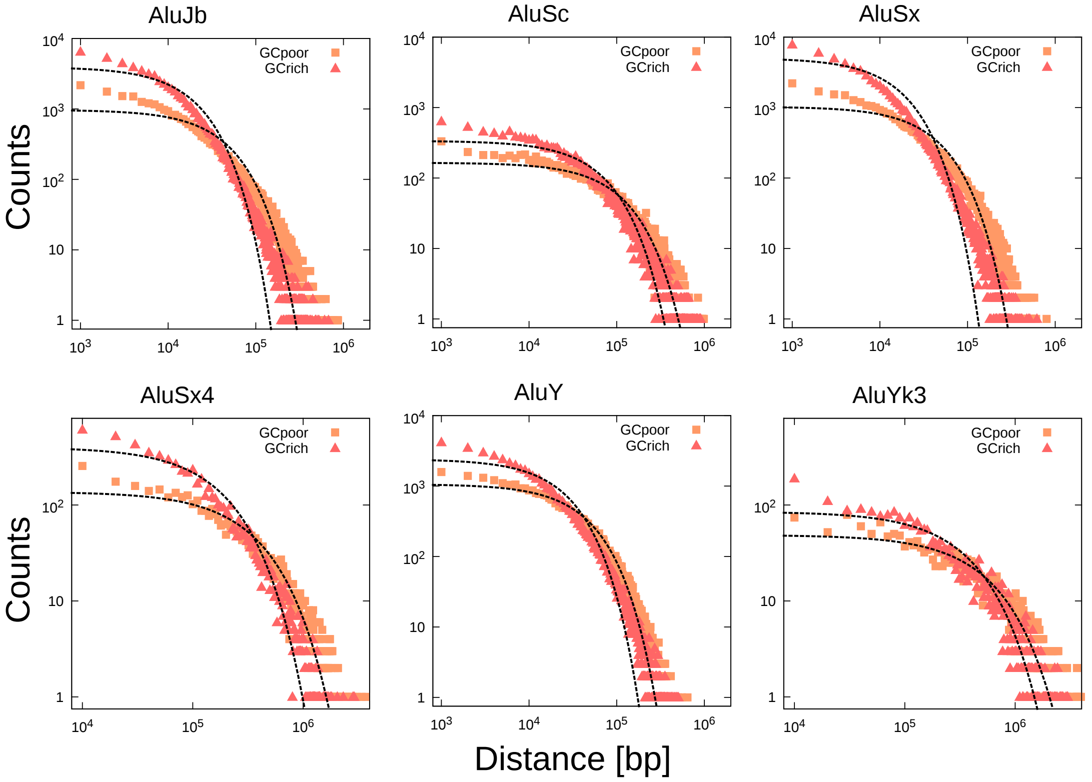

**Supplementary Figure S3. Examples of inter-Alus distance distributions in GC-rich and GC-poor regions.** Figure shows few examples of inter-Alus distance distribution for different subfamilies in GC-rich and GC-poor regions. Stick-breaking prediction for each distribution is shown (black dashed line).

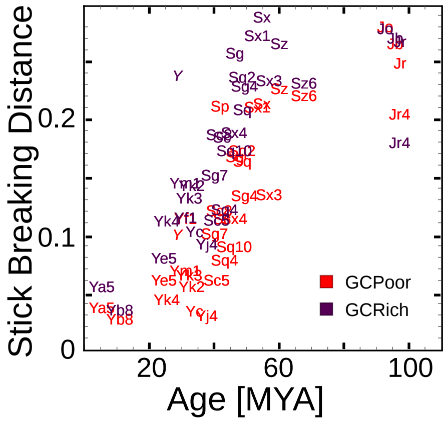

**Supplementary Figure S4.** Panel shows the distance from null model distribution as a function of the age for different Alu subfamilies in GC-rich and GC-poor regions.

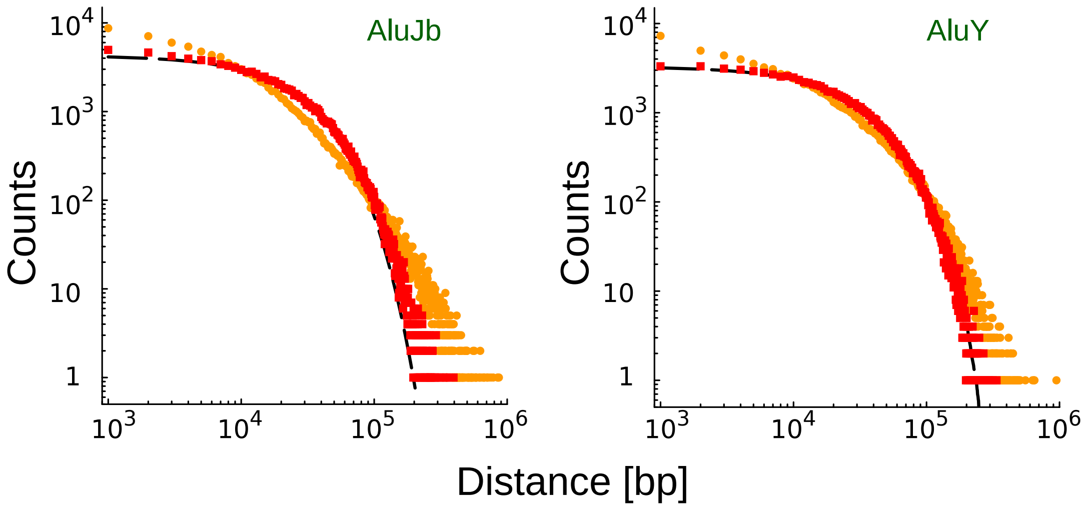

**Supplementary Figure S5. Effect of expansion on stick-breaking distribution.** We wanted to evaluate the effect of a "realistic" expansion due to TEs insertions and duplications on inter-REs distances. We calculated the size distribution of all the TEs with Jukes-Cantor divergence smaller than AluY and AluJb subfamily respectively. These TEs are likely to be younger than the Alus belonging to the two subfamilies, and could have contributed to expansion of inter-AluYs and inter-AluJbs distances. Data on genomic duplications in human larger than 1 kbp and identity > 90% (genome build GRCh37) were downloaded from *http://humanparalogy.gs.washington.edu/build37/build37.htm.* We selected those duplications that do not contain any insertion of AluY and AluJb in human genome (GRCh37). We simulated the random placement of a number of REs equal to the number of AluY and AluJb members present in human genome. Genome size was reduced to 2.5 Gbp and 2 Gbp for AluY and AluJb respectively in order to obtain for both the subfamilies a final $L_E \approx 2.8$ Gbp. Figure shows the actual distribution of distances for AluY and AluJb elements (orange dots) and simulated data (red square) of expanded distribution after random insertion of the selected TEs and duplicated regions. In the case of AluJb subfamily, duplicated regions are inserted twice, since data on duplications are likely to refer to events newer than AluJb insertion. Expanded distributions can not reproduce the empirical data and are well described by null model prediction (black dashed line). However, for AluJb we observed a small deviation of the tail from the null model distribution, possibly due to an effect similar to the one described by Sellis and coworkers Sellis et al. [2007]. This deviation is purely indicative, since the simulated expansion process does not reproduce a "real" expansion of human genome. Despite the many arbitrary choices we made, these results support the idea that a random expansion process is not sufficient to explain the empirical distributions.

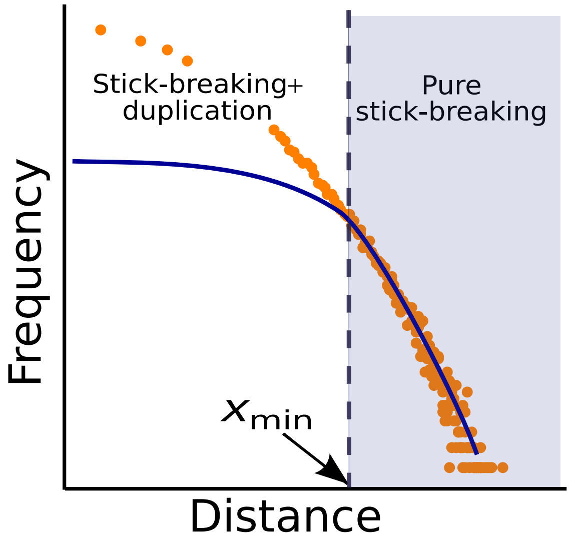

**Supplementary Figure S6. Schematic representation of source estimate procedure.** The inter-REs distance distribution (orange dots - data do not refer to any particular subfamily) for a value larger than a minimum value $x_{min}$ (black dashed line) is well approximated by a stick-breaking distribution (blue line). The same distribution at shorter distances is the sum of the stick-breaking and the source. The parameters of the stick-breaking distribution and the threshold $x_{min}$ can be optimized using a maximum likelihood approach.

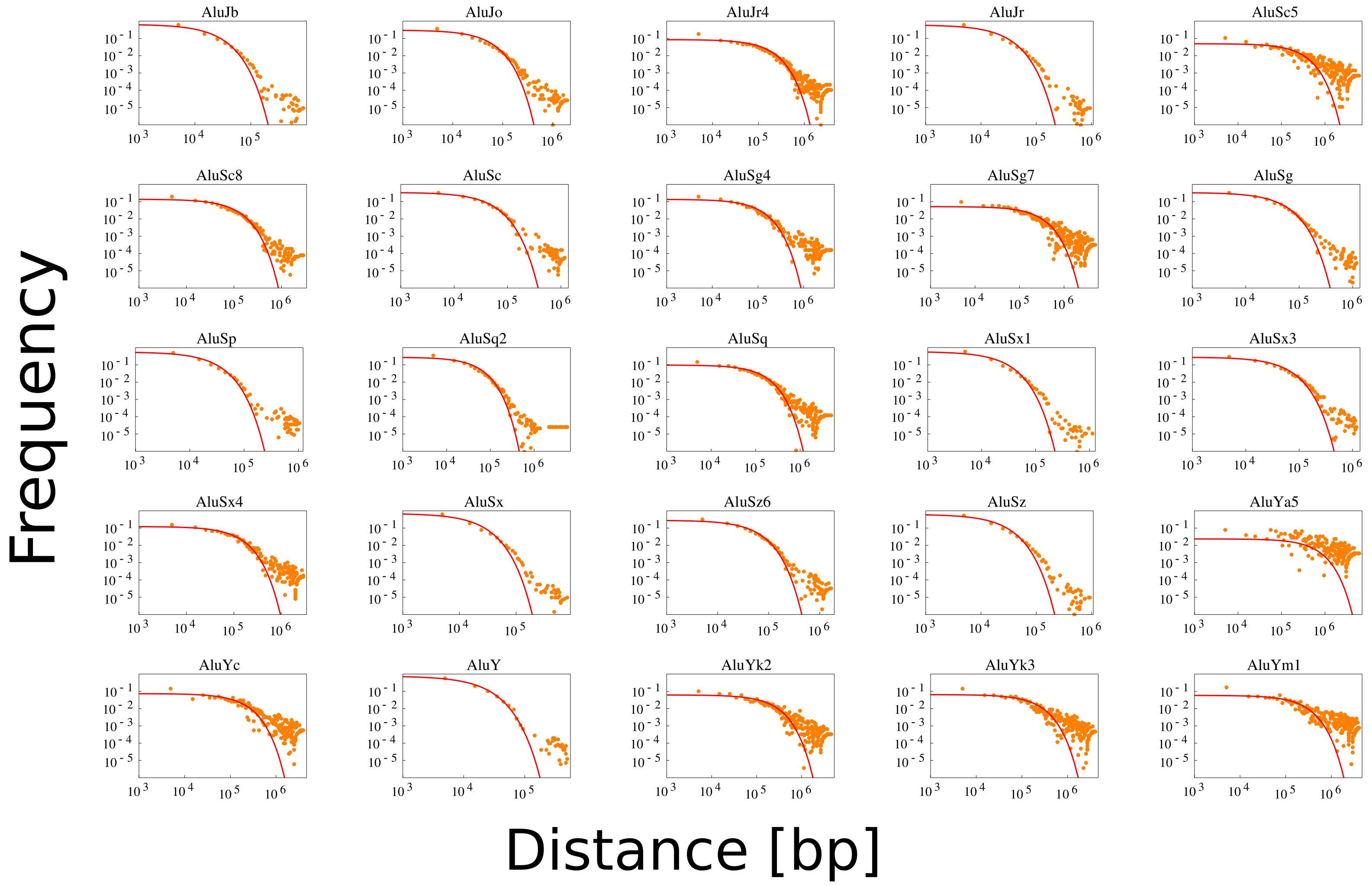

**Supplementary Figure S7. Estimated sources for Alu subfamilies.** The figure shows the result of source estimate procedure for 25 Alus subfamilies (orange dots). Most of the sources are well described by an exponential function, while older subfamilies are better described by a double exponential (continuous red lines). All the data obtained as difference between empirical distribution and optimized stick-breaking are shown, including distances larger than $x_{min}$ (in most of the cases $x_{min} \approx 10^5$). The exponential fit was performed using only data in the range $x < x_{min}$.

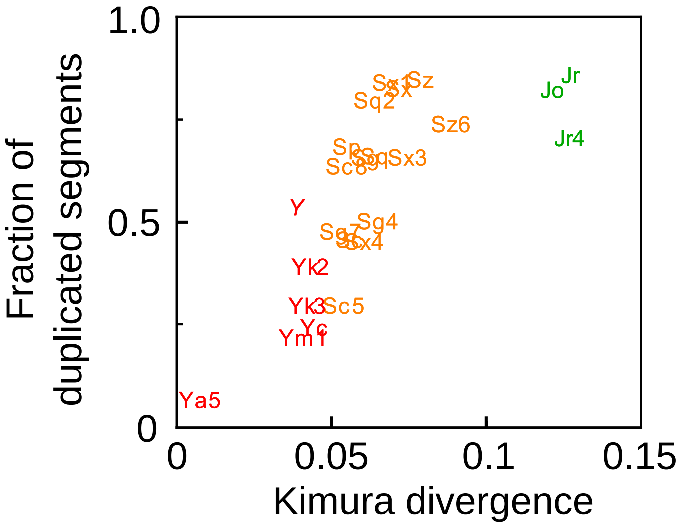

**Supplementary Figure S8.** The panel shows the fraction of duplicated segments in function of the Kimura divergence.
